# The PRC2-dependent epigenetic reprograming of the bladder epithelium exacerbates urinary tract infections

**DOI:** 10.1101/2020.10.22.350942

**Authors:** Chunming Guo, Mingyi Zhao, Xinbing Sui, Zarine Balsara, Songhui Zhai, Ping Zhu, Xue Li

**Author notes:** Equal first author (Chunming Guo, Mingyi Zhao). Equal second author (Xinbing Sui, Zarine Balsara). Correspondence (XL) or (XL).

## Abstract

Mucosal imprint sensitizes recurrent urinary tract infections (UTIs), a significant health and quality of life burden worldwide, which are associated with heightened inflammatory host response, severe basal cell hyperplasia and impaired superficial cell differentiation. Here, we show that bladder infections induce expression of *Ezh2*, the methyltransferase of polycomb repressor complex 2 (PRC2)-dependent epigenetic gene silencing program. In mouse models of UTIs, urothelium-specific inactivation of PRC2 reduces the urine bacteria burden. The mutants exhibit a blunted inflammatory response likely due to the diminished activity of *NF-κB* signaling pathway. PRC2 inactivation also improves urothelial differentiation and attenuates basal cell hyperplasia phenotype. Moreover, the Ezh2-specific small molecule inhibitors markedly improve disease outcomes of bladder superinfection and chronic cystitis. Taken together, these findings suggest that the UTI-induced epigenetic reprograming in the bladder urothelium likely contributes to the mucosal imprint, and further suggest that targeting PRC2 methyltransferase offers a non-antibiotic strategy to mitigate UTIs.

## Introduction

The ability to learn from the past experience offers an added advantage to defend against pathogens upon subsequent encounters. For example, a local infection in plants results in the epigenetic changes that lead to systemic acquired resistance when challenged by the same or unrelated pathogens(1). Invertebrates also display an enhanced response to secondary infection; such a memory-like feature persists and is transmissible to the next generation(2, 3). Vertebrates have a highly specialized adaptive immune system, which produces long-lasting immunological memory and enhanced responses to eliminate pathogens. Emerging evidence suggests that the innate immune cells also display a long-term enhanced inflammatory response, termed “trained immunity”(4), to challenge infections, suggesting that mammalian cells may have retained the ancestral mechanism of immunological memory to defend against infections(5, 6). The barrier epithelial cells are often the first to encounter pathogens. However, whether mammalian epithelial cells are able to retain the information of pathogen encounter and to respond accordingly during secondary infections have been poorly understood.

The bladder urothelium is a specialized epithelial barrier that prevents entry of irritants and pathogens in the urine. Unlike other epithelial barriers such as those of the skin and intestine, mature bladder urothelium has a very low proliferation index(7). The estimated urothelial cell turnover rate in rodents is once every 200 days. Nevertheless, bladder urothelium has a remarkable regenerative capacity to maintain its barrier function after injuries(8). For example, during urinary tract infections (UTIs), the quiescent urothelium becomes hyperproliferative and regenerates the superficial or umbrella cells within days to repair the damaged tissue(9–13). The rapid regenerative kinetics of the bladder urothelium appears to be biologically necessary for the restoration of the barrier function. Likewise, the exceedingly low turnover rate suggests that molecular changes of the urothelial cells could persist throughout the lifetime of bladder urothelium.

UTIs caused by uropathogenic *E. coli* (UPEC) are among the most common bacterial infections that afflict over 150 million people each year worldwide; and have the estimated 2-3 billion dollars of annual costs in the United States alone(14, 15). Even after appropriate antibiotic treatment, UTI patients have >25% chance of developing a recurrent infection in the following 6 months, suggesting that a prior history of bladder infection is an independent risk factor for subsequent UTIs(16–18). Intriguingly, bacteria induce bladder mucosal imprint, which sensitizes to recurrent and chronic diseases(19). Patients with chronic and recurrent UTIs display pathological mucosal remodeling including urothelial hyperplasia and a lack of terminal differentiation(20, 21). Analogous to trained immunity, mucosal imprint results in a heightened inflammatory response to bacterial infections(19). An exaggerated bladder inflammation, however, predisposes to symptomatic chronic and recurrent infections(22, 23). When mice are experimentally primed with either the bacterial or chemical injuries, these pretreated mice develop a heightened inflammatory host response but poor disease outcomes upon reinfections(24, 25). Rather than a self-limiting acute cystitis, the primed mice exhibit severe urothelial damage followed by chronic cystitis, pyelonephritis and mortality. Hence, the amplitude of bladder inflammatory response to bacterial infections is closely linked to clinical outcomes of UTIs from asymptomatic to life-threatening diseases(18).

Among the host defense strategies, exfoliation physically removes infected urothelial cells and intracellular bacterial communities (IBCs)(26–28), thereby facilitating bacterial clearance. However, this is a double-edged strategy as it damages the epithelial barrier and exposes the underlying tissue to pathogens and irritants, which could seed recurrent infections(29). In addition to exfoliation, the bladder urothelial cells are key in detecting UPEC through receptors including Uroplakin Ia (*Upk1a*) and Toll-like receptor 4 (*Tlr4*)(30–32), which activate downstream effectors such as the nuclear factor kappa B (*NF-κB*)-dependent inflammatory cytokines and chemokines. Along with other inflammatory mediators, the bladder urothelial cells produce a burst of interleukin 1a (*II-1α*), *Il-6* and *Il-8* to recruit immune cells such as neutrophils that are critical in actively clearing out the invading pathogens(31–33). Collectively, the bladder urothelium is the essential epithelial barrier and a central player in orchestrating the immunological responses to UTIs.

Exposure to pathogens may cause epigenetic remodeling of host cells(34–36). The epigenetic deposits such as histone H3 lysine 27 trimethylation (H3K27me3) have the potential to function as a molecular memory after pathogen clearance since they can persist through multiple cell cycles and several generations(37–39). H3K27me3 is tightly linked to chromatin compaction and transcription silencing(40–42). Intriguingly, exposure of human bladder urothelial carcinoma cells to UPEC *in vitro* induces expression of *Ezh2*, the methyltransferase of polycomb repressor complex 2 (PRC2)(43), which catalyzes formation of H3K27me3. We have previously reported that the PRC2-dependent H3K27me3 modification plays a critical role in the bladder urothelium during embryonic development(9). However, it’s potential role in mature urothelial cells during UTIs has not been reported yet. Here, we provide evidence suggesting that UPEC induces *Ezh2* in the urothelial cells; and the PRC2-dependent epigenetic program is tightly associated with an exaggerated inflammatory response and poor disease outcomes of UTIs. Moreover, we show that targeting the PRC2 activity, both genetically and pharmacologically, significantly mitigates severity of infections. These findings collectively suggest that the UTI-induced mucosal imprint is in part mediated by the PRC2-dependent epigenetic mechanism and further suggest that targeting the underlying epigenetic program offers an alternative non-antibiotic adjuvant strategy to ameliorate UTIs.

## Results

### Bacterial infection induces *Ezh2* gene expression

Wildtype C57Bl6 mice were infected by transurethral inoculation of human UPEC UTI89 directly into the bladder(44). To identify candidate UTI-induced epigenetic regulators, we performed whole transcriptome analysis of the bladder urothelium at one day post inoculation (dpi, Figure 1A). Among the differentially expressed genes (DEGs) between the UPEC-infected and vehicle controls, *Ezh2* was significantly upregulated (Table S1). Genes that encode the DEAD-box containing proteins Ddx26b and Ddx5, the SWI/SNF complex protein Smarcad1 and bromodomain protein Baz2b were also upregulated by UPEC infection. Several long intergenic noncoding RNAs including *Xist, Neat1, A330044P14Rik, Gm21781, Al480526* and *Gm26905* were also significantly induced. Other candidate epigenetic regulators including *Chst2, Uchl1, Padi1, Padi2* and *Inmt* were downregulated in the bladder urothelium after the infection.

**Figure 1.**
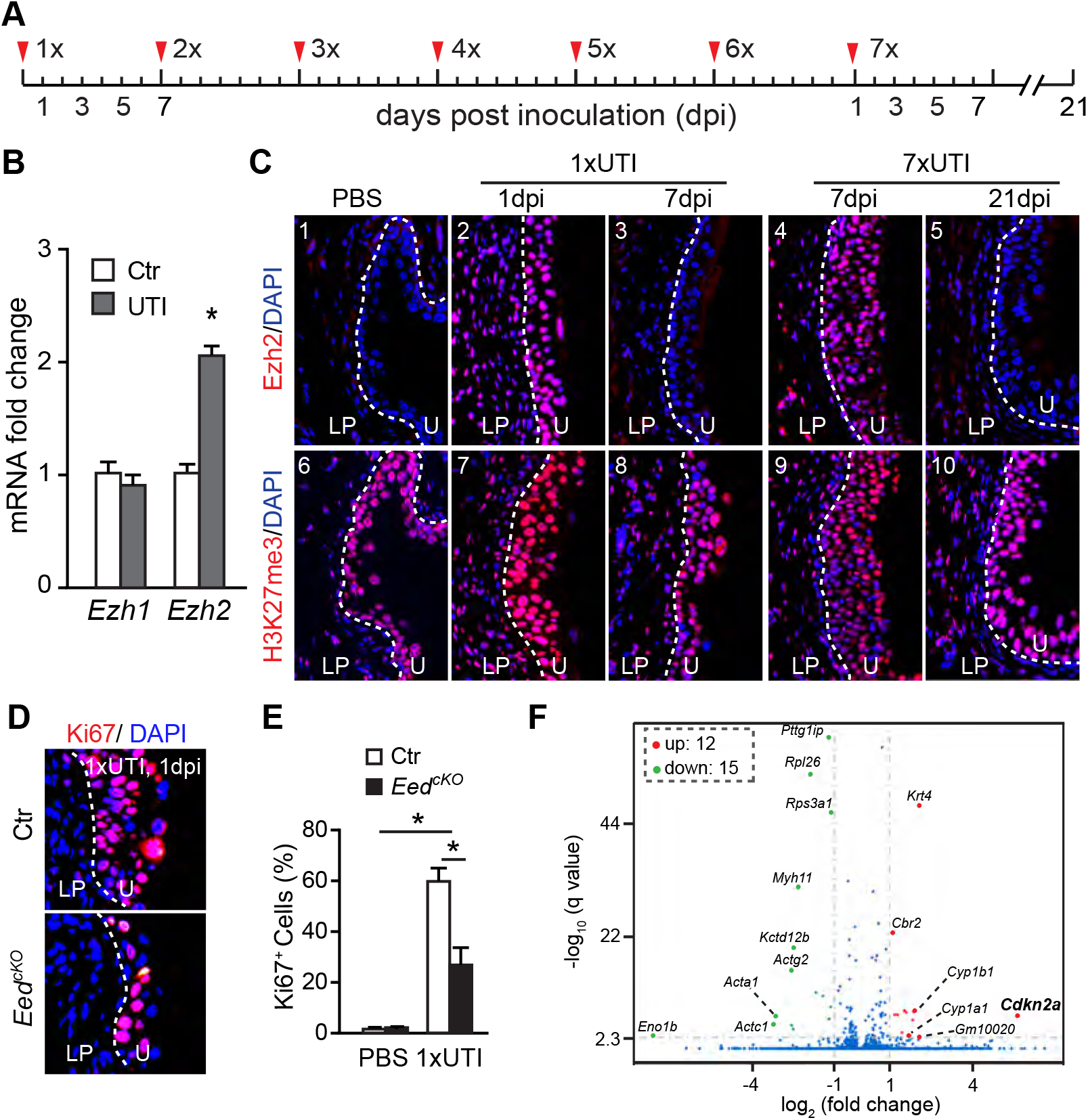
Bacterial infection induces *Ezh2* gene expression to promote hyper-proliferation of urothelial progenitor cells. (A) Schematic diagram of mouse models of acute (1x) and repeated (2x-7x) bladder infections. (B) Quantitative RT-PCR analyses of *Ezh1* and *Ezh2* mRNAs in the bladder urothelium at 24 hours after an acute infection. Control mice (Ctr) were treated with saline (PBS). *, *p*<0.05, Student’s *t*-test. (C) Ezh2 and H3K27me3 immunostaining (red) of mouse bladders that have been infected either acutely (1xUTI) or repeatedly (7xUTI). dpi, day-post-inoculation. Control mice were treated with PBS. Blue, nuclear DAPI counter staining; LP, lamina propria; U, urothelium; dashed line separates LP and U. (D and E) Ki67 immunostaining (red) of bladder sections of urothelium-specific *Eed* conditional knockout mice (*Eed*^cKO^) that are acutely infected and analyzed at 1dpi. Littermate heterozygous mice are used as controls (Ctr). Percentage of Ki67 positive (Ki67^+^) cells are quantified in D. *, *p*<0.05, Student’s *t*-test; PBS group, n=3; 1xUTI group, n=3 (Ctr) and 5 (*Eed^cKO^*). (F) Volcano blot of the differentially expressed genes (DEGs, *padj*<0.05, n=2) in adult *Eed^cKO^* urothelium.

We focused on the PRC2-dependent epigenetic repression program as it is known to play a critical role in urothelial differentiation during bladder development(9). Quantitative analysis confirmed that bacterial infection resulted in a significant increase of *Ezh2* (Figure 1B). Expression of other members of PRC2 complex including *Ezh1, Eed* and *Suz12* was not affected (Figures 1B and S1A). Ezh2 protein was barely detectable in the adult bladder urothelium by immunohistochemistry underlying physiological condition (Figure 1C, panel 1). In contrast, strong Ezh2 immunoreactivity was observed in the bladder urothelium and lamina propria layers at 1dpi (Figure 1C, panel 2). Consistently, strong H3K27me3 immunoreactivity was detected in the bladder urothelial and lamina propria cells (Figure 1C, panels 6-7). Similar to an acute infection, the repeated bladder infections also resulted in the markedly inductions of *Ezh2* and H3K27me3 in the bladder and moreover, *Ezh2* induction lasted for more than 7 days (Figure 1C, 7xUTI, panels 4-5 and 9-10). Together, these findings suggest that bladder infections may lead to the PRC2-dependent epigenetic remodeling by inducing expression *Ezh2* methyltransferase.

### Urothelium-specific *Eed* condition knockout blunts the proliferative response to UTI via the *p53* pathway

To examine a potential role of the PRC2-dependent epigenetic program in bladder urothelium during UTI, we used urothelium-specific *Eed* conditional knockout mice, *Eed^cKO^*, in which PRC2-dependent methyltransferase activity mediated by both *Ezh1* and *Ezh2* was eliminated from the bladder urothelium(9). An acute infection (1xUTI) resulted in the loss of cytokeratin 20-positive (Krt20^+^) superficial cells, accompanied by an increase in Krt5^+^ basal cells in both the mutants and controls (Figure S1B). Krt20^+^ superficial cells were recovered within 3-7 days in both *Eed^cKO^* and controls. While *Eed^cKO^* responded to the bladder infection by activating urothelial cell proliferation, the cell proliferation rate in *Eed^cKO^* mice was more than 2-fold less (<30%) than that of controls (~60%) based on Ki67 staining (Figures 1D and E). Similarly, ~15% of control and ~7% of *Eed^cKO^* urothelial cells were positively labeled by a mitotic marker phospho-histone H3 (Figure S1C and D, *p* < 0.05). Collectively, these findings suggest that urothelium-specific inactivation of PRC2 significantly blunts the hyper-proliferative response to bacterial infection.

To examine the PRC2-dependent transcriptional program in the developing and mature bladder urothelium, we profiled the urothelium-specific transcriptome using a Translating Ribosome Affinity Purification (TRAP) strategy(45). Urothelium TRAP RNAs were isolation from *Eed^cKO^* and controls at embryonic day 18.5 (E18.5) and adult bladders (Figure S1E). Transcriptomic analysis identified 197 differentially expressed genes (DEGs, q<0.05, n=2 per group) at E18.5 (Figure S1F). Among the DEGs, 159 of them were upregulated and 38 were downregulated. Consistent with the previous report(9), expression of cell cycle inhibitors including *Cdkn2a, Cdkn1a, En1, En2, Pax6* and *Six3* were aberrantly upregulated in the urothelium of *Eed^cKO^* mice (Figure S1F). Ingenuity Pathway Analysis (IPA) of the DEGs revealed that the *p53* molecular pathway was significantly activated (Figure S1G). Among *p53* gene targets, *Cdkn1a, Bbc3, Trp53inp1* and *TP73* were highly upregulated. Additional pathways including the Aryl hydrocarbon receptor, G1/S checkpoint and glioblastoma multiforme signaling pathways were upregulated while the cyclins and cell cycle regulation pathway was significantly downregulated. Similar to the developing urothelium, *Cdkn2a* was also upregulated in adult urothelium of *Eed^cKO^* mice (Figure 1F). A total of 27 DEGs was observed from the adult urothelium, which was much fewer than that from the embryonic urothelium, consistent with the observation that the adult *Eed* knockout bladders were histologically indistinguishable from littermate controls. Taken together, results from these transcriptomic analyses suggest that the PRC2-dependent epigenetic program promotes urothelial cell proliferation possibly via the *p53* tumor suppressor pathway.

### *PRC2* heightens the inflammatory response to bladder infections

Bladder infections resulted in the immune cell infiltration, which was observed in all three layers of the bladder including the urothelial, lamina propria and smooth muscle layers (Figure 2A). After an acute bacterial infection, damage to the urothelium as indicated by the loss of Krt20^*+*^ superficial cells and increase of Krt5^*+*^ basal cells were apparent and comparable between *Eed^cKO^* knockouts and controls (Figure S1B). IBCs formed at 3hpi or 6hpi were also comparable between the knockouts and controls (Figures S2A). Interestingly, the urine bacterial counts were lower in the knockouts (Figure 2C). *Eed^cKO^* bladders appeared to have fewer infiltrating immune cells when compared to controls (Figure 2A). To quantify the difference, we analyzed bladder inflammation using the criteria based on immune cell infiltration, edema, location of neutrophils infiltration and loss of urothelium(46). The overall inflammatory score of *Eed* knockouts was approximately 2-fold less when compared to controls (Figure 2B). At 2 hour-post-inoculation (hpi), acute infection resulted in a rapid upregulation of inflammatory regulators including *Il-6*, suppressor of cytokine signaling-3 (*Socs3*), C-C motif chemokine ligand 2 (*Ccl2*)/monocyte chemoattractant protein 1 (*MCP1*), C-X-C motif chemokine ligand 1 (*Cxcl1*)/neutrophil-activating protein 3 (*NAP-3*)/(*KC*) and chemokine (C-X-C motif) receptor 2 (*Cxcr2*) in the bladder of both *Eed* knockouts and controls. Consistent with the observation that *Eed^cKO^* mice had lower inflammatory score, expression of *Il-6, Socs3, Ccl2, Cxcr2* and chemokine (C-C motif) receptor 2 (*Ccr2*) and chemokine (C-X-C motif) receptor 2 (*Cxcr2*) were significantly lower in the knockouts than in controls at 24hpi (Figure 2D). Expression of *Cxcl1* was significantly lower at both 2hpi and 24hpi in the mutants.

**Figure 2.**
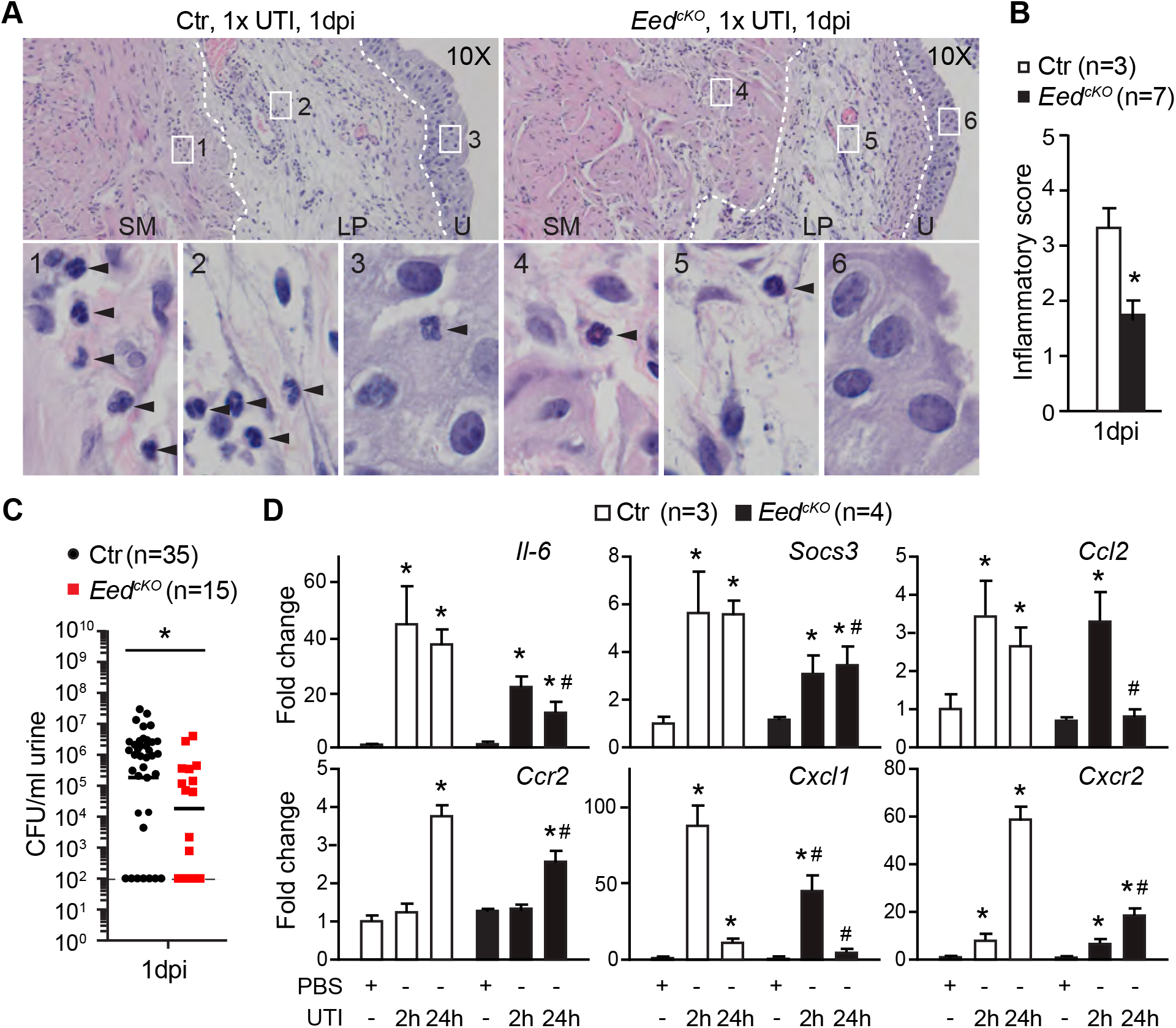
Genetic inactivation of PRC2 significantly blunts inflammatory host response to bladder infections. (A) Representative histological images of *Eed^cKO^* bladders one day post-inoculation (1xUTI and 1dpi) of UTI89 into the bladder. Control (Ctr) mice were heterozygous littermates. Arrowheads, polymorphonuclear leukocytes; LP, lamina propria; SM, smooth muscle; U, urothelium; dashed lines separate SM, LP and U. (B) Inflammatory scores based on histological data as shown in A. *, *p*<0.05, Student’s *t*-test. (C) Urine bacterial counts at 1dpi. \**p*<0.05, Mann-Whitney test. (D) Quantitative RT-PCR analyses of expression of inflammatory markers in the bladder of *Eed^cKO^* mice at 2 hours and 24 hours post-inoculation. Heterozygous littermates were used as controls (Ctr). *Gapdh* was used as an internal control. *, *p*<0.05, vs PBS with the same genotype; #, *p*<0.05, vs control (Ctr) with the same treatments; Student’s *t*-test.

Bladder infection results in a rapid nuclear accumulation of *NF-**κ**B* in superficial cells and, subsequently, in intermediate and basal cells(30). *NF-**κ**B* is a central transcription regulator that activates expression of a number of gene targets including cytokines and chemokines(47). Inhibitor of **κ**B (*I**κ**B*) binds and sequesters *NF-**κ**B* in the cytoplasm, thereby inhibiting the *NF-**κ**B*-dependent inflammatory response(48). We found that expression of *I**κ**Bα* was significantly upregulated in the bladder urothelium of *Eed* knockout mice than in controls (Figure 3A). Consistently, anti-I**κ**Bα immunoreactivity was markedly stronger in *Eed^cKO^* mice compared to controls (Figure 3B). Chromatin immunoprecipitation (ChIP) further showed that the occupancy of a repression histone mark H3K27me3 was significantly reduced, while the epigenetic activation mark H3K4me3 was significantly enriched at the *IκBa* promoter region of *Eed^cKO^* mice (Figure 3C). Furthermore, while many superficial cells with strong nuclear *NF-**κ**B* staining were observed in controls, fewer were found in *Eed^cKO^* mice (Figure 3D). Taken together, these observations suggest that the crosstalk between PRC2 and *NF-κB* molecular pathways promotes the inflammatory host response during bladder infections.

**Figure 3.**
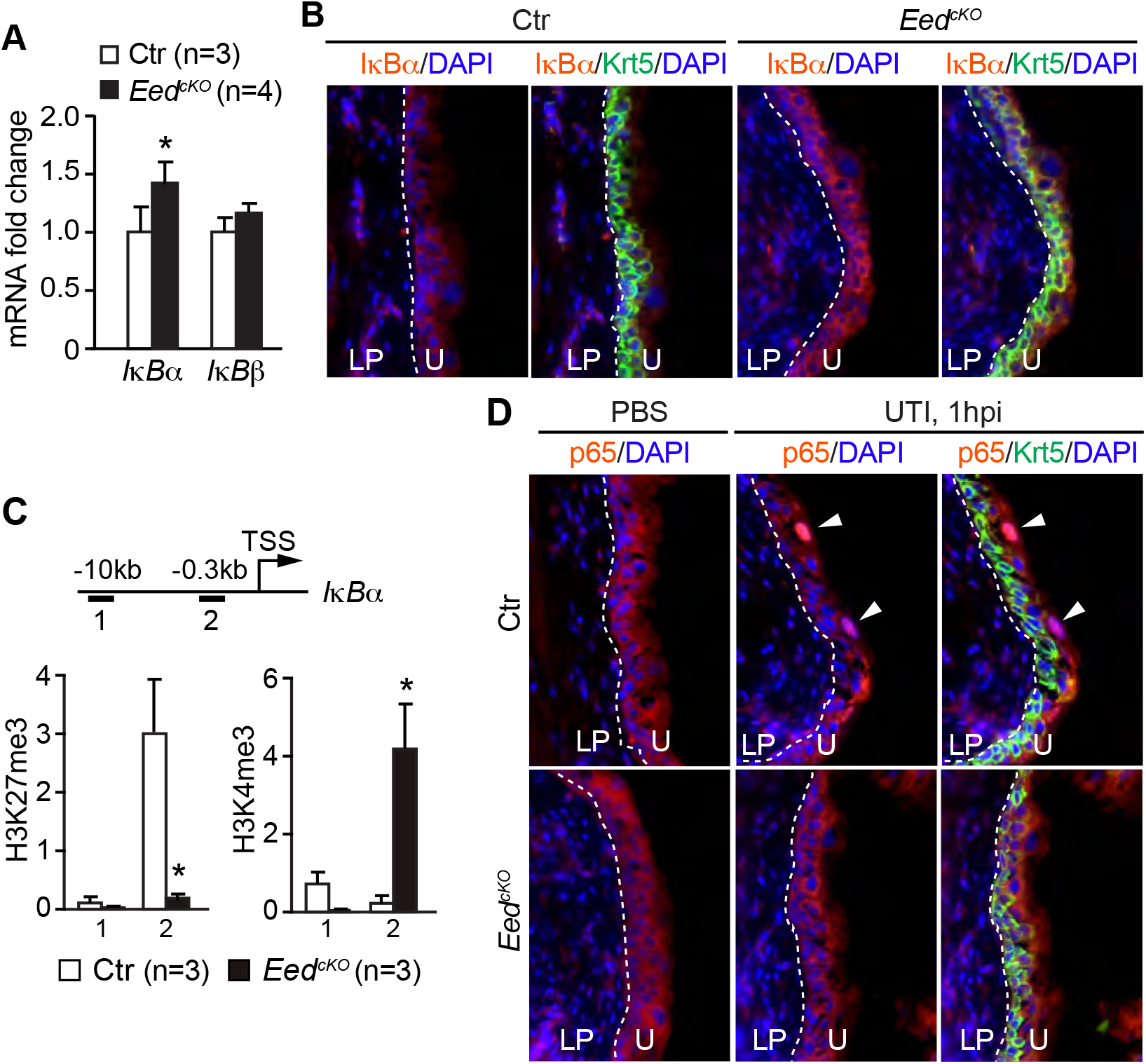
PRC2 enhances activity of the *NF-κB* signaling pathway in the bladder urothelium during acute bacterial infection. (A) Quantitative analysis of *I**κ**Bα* and *I**κ**Bβ* mRNAs in the bladder urothelium of control and *Eed^cKO^* mice without infection. *Gapdh* was used as an internal control. *, *p*<0.05, Student’s *t*-test. (B) Immunostaining of I**κ**Bα (red) and Krt5 (green) of bladder sections from *Eed^cKO^* mice without bacterial infection. (C) Chromatin immunoprecipitation (ChIP) assay of *I**κ**Bα* locus (schema) using H3K27me3- and H3K4me3-specific antibodies. TSS, transcription start site. *, *p*<0.05, Student’s *t*-test. (D) Immunostaining of NF-**κ**B p65 subunit (red) and Krt5 (red) of bladder sections from *Eed^cKO^* mice at 1 hour post inoculation (hpi). White arrowheads, nuclear localized p65. Ctr, heterozygous littermates; PBS, uninfected mice; Green, Krt5^+^ basal cells; blue, DAPI counter staining; LP, lamina propria; U, urothelium; dashed lines separate U from LP.

### A history of UTI heightens the inflammatory host response

To examine the impact of a prior history of bladder infections, mice were inoculated repeatedly for up to 7 times with a 7-day convalescence after each infection (Figure 4A). Mice developed bacteriuria after the first infection with approximately 10^6^ colony forming units (CFU) of bacteria in the urine at 1dpi (Figure 4B, 1xUTI, 1dpi, Ctr). The second infection (2xUTI) resulted in a significantly lower urine bacterial count (~10^3^ CFU, p<0.05). However, additional reinfections (3xUTI-7xUTI) resulted no further decrease in bacteriuria, similar to previous reports using different UPEC and mouse strains(49–51). unlike the controls that were inoculated repeatedly with saline, histological analyses showed that repeated infections increased the inflammatory score and caused severe urothelial hyperplasia of the bladder (Figures 4C-E and S2B, Ctr). Expression of *Krt20* and Uroplakin 3a (*Upk3a*) were undetectable at 1, 3 and 7dpi after repeated infections (Figures 4E and S2C, Ctr). The Krt5^+^ basal cells were expanded and occupied the entire thickness of the bladder urothelium, strongly suggesting that the repeated infections caused exfoliation of both intermediate and superficial cells. Collectively, these findings suggest that a history of UTI heightens the inflammatory host response to reinfections and causes pathological remodeling of the bladder urothelium, including basal cell hyperplasia and a lack of terminal superficial cell differentiation.

**Figure 4.**
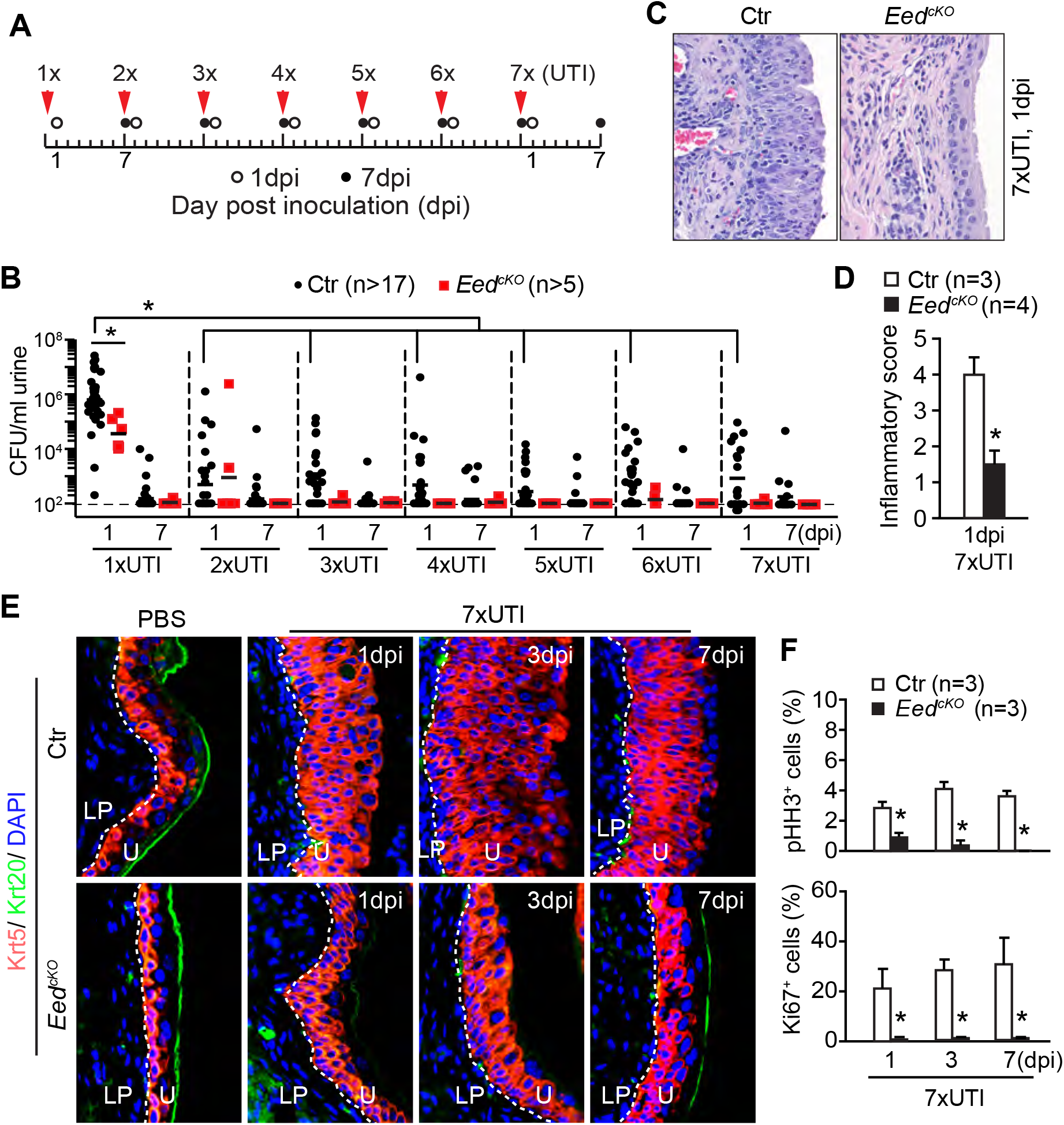
Genetic inactivation of PRC2 in the bladder urothelium attenuates inflammatory host response to the repeated bladder infections. (A) Schematic diagram of repeated infections. (B) Urine bacterial counts during repeated infections. \**p*<0.05, Mann-Whitney test. (C) Representative images of histological staining of bladder tissues from *Eed^cKO^* mice received 7xUTI at 1 day-post-inoculation (dpi). Ctr, littermate heterozygous control. (D) Inflammatory scores based on histological data as shown in C. \**p*<0.05, Student’s *t*-test. (E) Immunostaining of Krt5 (red) and Krt20 (green) of bladder sections from *Eed^cKO^* mice after 7xUTI. Blue, DAPI counter staining; LP, lamina propria; U, urothelium; dashed lines separate U from LP. (F) Analysis of pHH3^+^ and Ki67^+^ cells in the bladder urothelium of *Eed^cKO^* and control mice after 7xUTI. \**p*<0.05, Student’s *t*-test.

### *Eed* deletion improves outcomes for both acute and repeated infections

Compared to controls, urinalysis showed that the bacterial counts were significantly lower in *Eed^cKO^* mice at 1dpi after 1xUTI (Figure 4B). The same trend persisted during repeated infections (2xUTI to 7xUTI). Urine bacteria were reduced to almost undetectable in *Eed^cKO^* mice at 1dpi and 7dpi during reinfections (2-7x UTIs). *Eed* knockout bladders had significantly fewer infiltrating immune cells, less edema and milder urothelial hyperplasia when compared with controls (Figures 4C-E and S2B-D). The overall inflammatory scores were also lower in the knockouts than controls (Figure 4D). Urothelial cell proliferation, as indicated by both pHH3 and Ki67 markers, was significantly attenuated in *Eed^cKO^* mice compared to controls (Figures 4F, S2C and D). Krt20 immunoreactivity was not detected at 1dpi and 3dpi in *Eed* knockouts (Figure 4E), indicating a complete superficial cell exfoliation after repeated infections. However, abundant Krt20^+^ superficial cells were observed at 7dpi in the knockouts but not in controls (Figure 4E). Consistently, abundant Upk3a^+^ intermediate and superficial cells were observed in *Eed^cKO^* mice but not in controls (Figure S2C). Thus, genetic inactivation of PRC2 in the bladder urothelium improves outcomes of both the acute and repeated bladder infections.

### *Eed* deletion improves outcome of the C^+^UTI superinfection

Mice primed with cyclophosphamide (CPP) one day before the acute infection, the CPP plus UTI (C^+^UTI) superinfection(24), have severe outcomes including high levels of urine bacterial counts that persisted for more than four weeks (Figures 5A and S3A). The superinfected bladders had apparent edema, urothelial damage and basal cell hyperplasia with an inflammatory score of 5 (Figures 5B and C). *Krt20* and *Upk3a* were undetectable at 1dpi and the urothelium consisted entirely of Krt5^+^ basal cells (Figure 5D and S3B). While *Upk3a* expression began to recover at 7dpi after the C^+^UTI superinfection, expression of Krt20 was not detected before 14dpi in controls. High degrees of basal cell proliferation were observed at all stages analyzed (Figures S3C and D).

**Figure 5.**
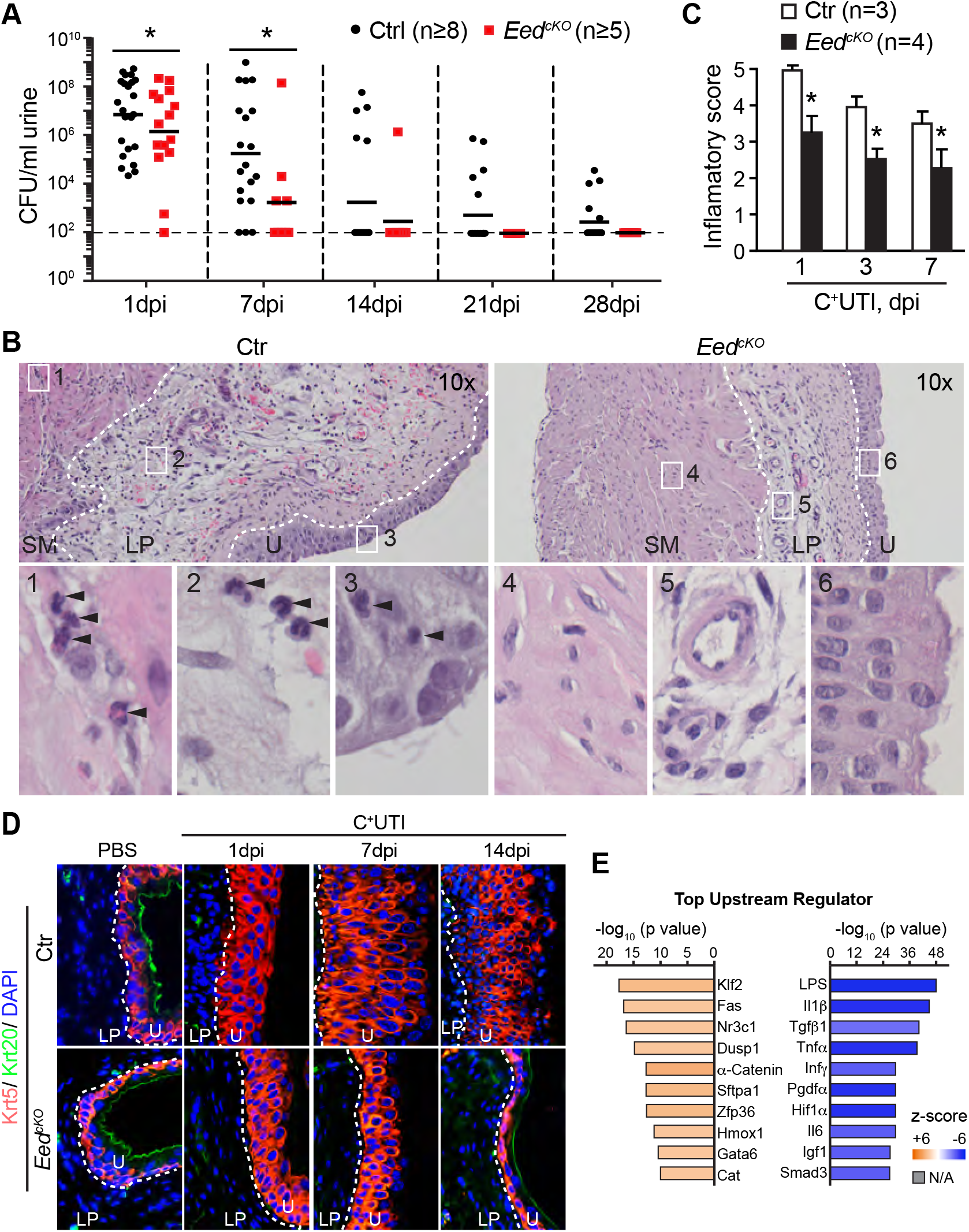
Inactivation of PRC2 in the bladder urothelium attenuates inflammatory host response to the C^+^UTI superinfection. (A) Urine bacterial counts of mice after the C^+^UTI superinfection. \**p*<0.05, Mann-Whitney test, dpi, day-post-inoculation. (B) Representative histological images of bladder sections from *Eed^cKO^* mice one day after the C^+^UTI superinfection. Ctr, littermate heterozygous mice; LP, lamina propria; SM, smooth muscle; U, urothelium; dashed lines separate SM, LP and U; arrowheads, polymorphonuclear leukocytes. (C) Inflammatory scores based on histological data as shown in B. \**p*<0.05, Student’s *t*-test. (D) Immunostaining of Krt5 (red) and Upk3a (green) of bladder sections from *Eed^cKO^* mice after the C^+^UTI superinfection. Blue, DAPI counter staining; LP, lamina propria; U, urothelium; dashed lines separate U from LP. (E) The top predicted Ingenuity Pathway Analysis (IPA) upstream regulators of the differentially expressed genes (*padj*<0.05, n=2) identified from the bladder urothelium of *Eed^cKO^* mice at 1dpi after the C^+^UTI superinfection.

Compared to controls, the *Eed* knockout mice had significantly lower urine bacterial burden at 1dpi; and bacteriuria was largely cleared within two weeks of the C^+^UTI superinfection (Figure 5A). Inflammatory score was also lower in *Eed^cKO^* mice (Figure 5B and C). Exfoliation of *Krt20^+^* superficial cells was observed in both the knockouts and controls at 1dpi, implying that the knockout mice had sustained a similar level of the initial urothelial damage. However, two weeks later at 14dpi, *Krt20* expression began to recover in the knockouts but not in controls (Figure 5D). Similarly, *Upk3a* expression was absent from controls but was clearly detected in the knockouts (Figure S3B). Basal cell hyperplasia was milder in the knockouts, which was coupled with a significantly lower cell proliferation rate at all stages examined from 1dpi to 14dpi (Figures S3C and D). Together, these findings suggest that conditional knockout of *Eed* in the bladder urothelium dampens the inflammatory host response while improving bacterial clearance and urothelial regeneration in bladder superinfection.

To better understand the PRC2-dependent inflammatory program, we performed whole genome transcriptomic analysis of the urothelium of superinfected bladders at 1dpi. We found that 124 genes were significantly downregulated and 62 genes were significantly upregulated in *Eed^cKO^* mice in comparison with controls (Figure S3E). Among the DEGs, a list of inflammatory markers including *Cxcl2, Cxcl3, Ccl2, Ccl3, Il1β, Il11* and *Ptgs2* (aka *Cox2*) were significantly downregulated in *Eed^cKO^* mice (Figure S3E), consistent with the findings that the inflammatory score was lower. Furthermore, IPA analysis revealed that molecular pathways such as the acute phase inflammation and *Il-8* neutrophil chemotaxis were significantly reduced (Figure S3F). Among the top predicted upstream regulators, induces such as lipopolysaccharides (LPS), *Il-1β*, Tumor necrosis factor alpha (*Tnfα*), Interferon gamma (*Infg*) and *Il-6* were significantly suppressed while inflammatory modulators including Kruppel-like factor 2 (*KLF2*)(52), *Nr3c1* (aka. Glucocorticoid receptor)(22), Zinc finger protein 36 (*Zfp36*)(53) and Heme Oxygenase 1 (Hmox1)(54) were significantly enhanced. Particularly noteworthy were suppression of *Cox2* and activation of glucocorticoid molecular pathways (Figure 5E); mice treated with either Cox2 inhibitor or glucocorticoid agonist significantly reduces chronic cystitis(22, 23). Collectively, these findings strongly suggest that the PRC2-dependent epigenetic program in bladder urothelium exacerbates the inflammatory host response.

### Small molecule Ezh2 inhibitors mitigate the C^+^UTI superinfection

To further explore the role of *Ezh2* in bladder infection, we examined the potential beneficial effects of Ezh2 small molecule inhibitors DZNep and EPZ005687, both of which reduce H3K27me3 and tumor growth *in vivo*(55). Because of the long half-life of H3K27me3 but short half-life of the inhibitors(39, 56), mice were pretreated with three doses of the inhibitors to deplete H3K27me3 before bacterial inoculation and 2 extra doses post-inoculation (Figure 6A). DZNep, a potent inhibitor of methyltransferases including *Ezh2*(56), markedly reduced H3K27me3 in the bladder (Figure 6B). DZNep treatment resulted in a mild but significant reduction of urine bacteria compared with the vehicle treatment in the C^+^UTI superinfection (Figure 6C). Bacterial counts in the bladder and kidneys were not affected by DZNep treatment (Figure S4A). Unlike the vehicle controls, the DZNep-treated mice displayed fewer infiltrating neutrophils, mild basal hyperplasia, blunted expression of *Ezh2*, and increased expression of *Upk3a* and *Krt20* (Figures 6D-G and S4B). EPZ005687 is an *Ezh2*-specific inhibitor(57), which did not affect UPEC growth or variability *in vitro*, nor the bacterial counts in the bladder or kidneys (Figure S4C-E). It did, however, decrease modestly bacteriuria at 3dpi (Figure S4D). EPZ005687 treatment also decreased the *NF-**κ**B* pathway activity as indicated by fewer cells with nuclear p65 staining (Figure S4F) and a reduction of the inflammatory score (Figures 6H and S4G). Expression of the urothelial cell differentiation marker *Upk3a* was also preserved (Figure 6I), and concordantly urothelial proliferation was significantly reduced in the EPZ005687-treated mice (Figure 6J). Collectively, these findings suggest that *Ezh2* inhibitors ameliorate effects of bladder superinfection.

**Figure 6.**
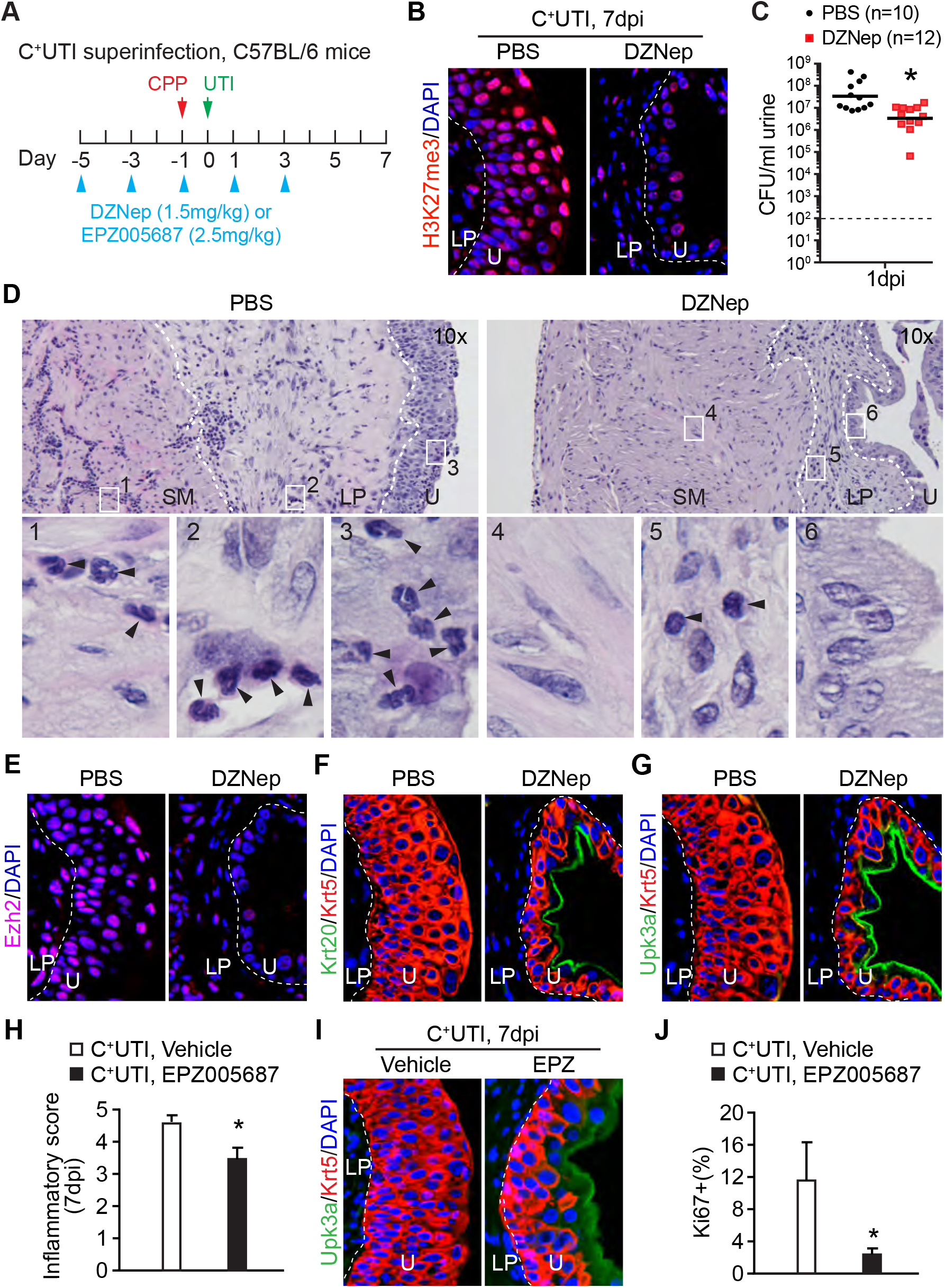
Ezh2 small molecule inhibitors blunt inflammatory host response and improve outcomes of the C^+^UTI superinfections. (A) Schematic diagram of the experimental regimen using C57Bl/6 mice. (B) H3K27me3 immunostaining in the bladder urothelium of mice treated with DZNep. PBS, vehicle control; blue, DAPI nuclear counter staining. LP, lamina propria; U, urothelium; dashed lines separate U from LP. (C) Urine bacterial counts at 1 dpi. \**p*<0.05, Mann-Whitney test. (D) Representative histological images of bladder sections from C57Bl/6J mice 7 days after the superinfections. SM, smooth muscle; arrowheads, polymorphonuclear leukocytes. (E-G) Immunostaining of Ezh2 (E, purple), Krt20 and Krt5 (F, green and red, respectively) and Upk3a and Krt5 (G, red and green, respectively) of the bladder urothelium of superinfected mice at 7dpi. (H) Inflammatory scores of the superinfected mice treated with Ezh2-specific inhibitor EPZ005687. \**p*<0.05, Student’s *t*-test, n=5. (I) Immunostaining of Upk3a (red) and Krt5 (green) of the bladder urothelium of superinfected mice at 7dpi. EPZ, EPZ005687; blue, DAPI nuclear counter staining. (J) Percentage of Ki67^+^ proliferating cells in the bladder urothelium of the EPZ005687-treated and super-infected mice. \**p*<0.05, Student’s *t*-test.

### Ezh2 small molecule inhibitors improve outcomes of chronic cystitis

Unlike C57Bl6 mice, C3H/HeN mice are susceptible to persistent bacteriuria or chronic cystitis(19, 22, 23). We therefore used this mouse strain to examine whether Ezh2 inhibitors may decrease the risk of chronic infection (Figure 7A). DZNep reduced H3K27me3 levels in the bladder of wildtype C3H/HeN mice (Figure 7B). However, it did not have major impact on the urine bacterial counts in C3H/HeN mice (Figure S5A). Nevertheless, neutrophil infiltration and inflammatory score were significantly lower in the DZNep-treated mice (Figures 7C and D). Apparent bladder edema, neutrophil infiltration and urothelial hyperplasia were found among the vehicle-treated controls at 10dpi (Figure 7C). These histological observations, however, were rarely detected in DZNep-treated mice. Consistently, high levels of *Upk3a* expression were detected in DZNep-treated mice but not vehicle-treated controls (Figure 7E). Similar to DZNep, EPZ005687 significantly blunted the induction of *Ezh2*, reduced H3K27me3 levels and decreased the overall inflammatory scores in C3H/HeN mice (Figures 7F-I and S5D). These mice also exhibited a mild basal cell hyperplasia phenotype with a concordant increase of terminal differentiation marker *Upk3a*. EPZ005687 alone had minimal effects on the bacterial counts in the urine, bladder and kidneys, although bacteriuria was moderately lower at 1dpi (Figures S5B and C). Taken together, these findings suggest that *Ezh2* inhibitors decrease inflammation and increase urothelial cell differentiation in chronic cystitis.

**Figure 7.**
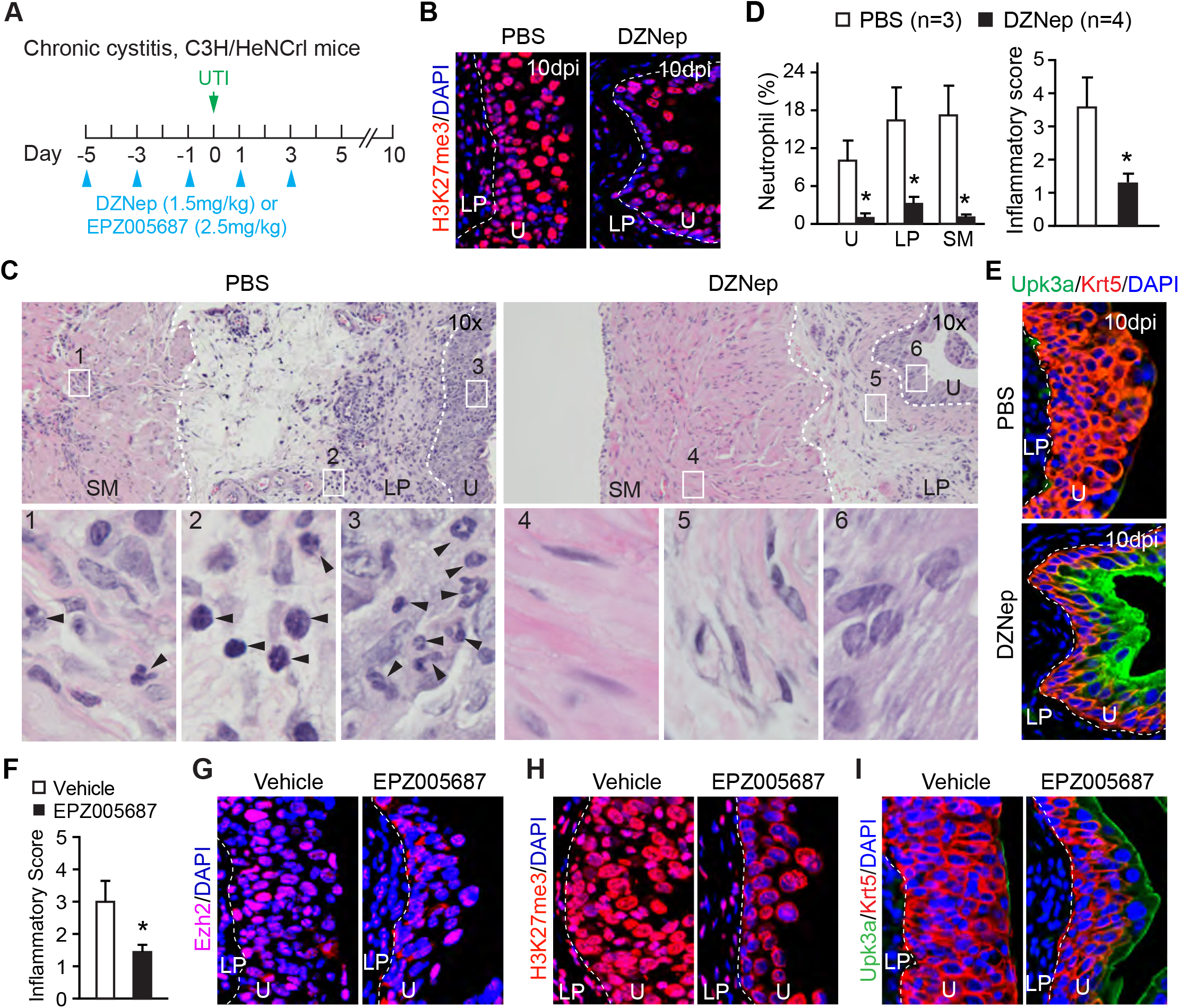
Ezh2 small molecule inhibitors blunt inflammatory host response and improve outcomes of the chronic cystitis. (A) Schematic diagram of the experimental regimen using C3H/HeN mice. (B) H3K27me3 immunostaining in the bladder urothelium of mice at 10dpi. Blue, DAPI nuclear counter staining; LP, lamina propria; U, urothelium; dashed lines separate U from LP. (C) Representative histological images of bladder tissues from C3H/HeN mice at 10dpi. SM, smooth muscle. Arrowheads, polymorphonuclear leukocytes. (D) Neutrophil infiltration and inflammatory score based on histological data as shown in C. \**p*<0.05, Student’s *t*-test. (E) Krt5 (red) and Upk3a (green) immunostaining of the bladder sections at 10dpi. (F) Inflammatory score of vehicle (n=3) or EPZ005687-treated (n=3) C3H/HeN mice at 7dpi. \**p*<0.05, Student’s *t*-test. (G-I) Immunostaining of Ezh2 (G, purple), H3K27me3 (H, red) and Upk3a and Krt5 (I, green and red, respectively) of the bladder urothelium of mice at 7dpi.

## Discussion

This study uncovers a previously unknown role of the PRC2-dependent epigenetic repression program in regulating the inflammatory host response to bacterial infections in the bladder. Findings here suggest that the bacteria-induced epigenetic remodeling of the bladder urothelium exacerbates inflammation and urothelial injury, thereby leading to poor outcomes of bladder infections. Collectively, these findings highlight the possibility of epigenetic basis of mucosal imprint and further suggest that targeting the underlying PRC2-dependent epigenetic program offers a non-antibiotic strategy to ameliorate UTIs.

UPEC induce expression of a list of epigenetic regulators in the bladder. Among them, *Ezh2* is significantly upregulated in both the bladder urothelium and underlying mesenchymal cells. By conditionally deleting *Eed* in the bladder urothelium, a strategy to inactivate PRC2 methyltransferase activity mediated by both *Ezh1* and *Ezh2*, our findings strongly suggest that the PRC2-dependent epigenetic program in the urothelial cells plays a critical role in facilitating the heightened inflammatory host response to bladder infections. *Ezh2* induction is tightly linked to the increased expression of inflammatory cytokines and chemokine, severe basal cell hyperplasia and impairment of urothelial cell terminal differentiation. While the underlying mechanism of PRC2-mediated inflammatory response in the bladder remains to be defined, our findings suggest that it promotes the NF-**κ**B signaling pathway by epigenetic repression of the *IκBa* locus. Indeed, the UTI-induced NF-**κ**B activation, as indicated by NF-**κ**B nuclear localization, is markedly reduced in *Eed^cKO^* mice. In addition to the PRC2-dependent histone methylation, *Ezh2* may also function in the methyltransferase-independent manner to enhance *NF-**κ**B* activity(58). It is worth noting that *NF-**κ**B* directly regulates *Ezh2* expression in human fibroblasts(59), suggesting a feedback regulation between *Ezh2* and *NF-**κ**B* molecular pathways may exaggerate and extend inflammatory host response to bladder infections. Importantly, the small molecule Ezh2 inhibitors significantly lessened bladder inflammatory responses to the acute, chronic and super infections. The treated mice have improved outcomes to bladder infections including decreased bacteriuria and enhanced urothelial regeneration. While the design of this study could not differentiate the prophylactic and therapeutic effects, these findings nevertheless provide the proof of concept evidence that targeting the PRC2-dependent epigenetic program offers a non-antibiotic adjuvant strategy to ameliorate UTIs.

Epigenetic activation has been linked to the trained immunity-like inflammatory memory in epithelial cells including the skin and airway epithelial progenitors(60, 61). However, epigenetic repression in modulating inflammatory host response to bacterial infections in the bladder has not been reported until now. Given the quiescent nature of bladder urothelium and how the PRC-dependent H3K27me3 modification persists through cell cycles and generations, we speculate that the PRC2-dependent epigenetic reprograming will have a lasting impact on the pathophysiology of the bladder urothelium. The virulence factor a-hemolysin is required in attenuating host Akt/protein kinase B signaling pathway during UPEC infection (62) and, at the a-hemolysin-dependent manner, UPEC also abrogate global histone H3 and H4 acetylation in Sertoli cells to promoter cell survival(63). In addition to a-hemolysin, other pore-forming bacterial toxins have been reported to mediate dephosphorylation of histone H3 and deacetylation of histone H4(34). Hultgren and colleagues have previously shown that a primary bladder infection leaves behind a molecular imprint on the bladder tissue that lasts for more than seven months and, importantly, sensitize recurrent infections(19). Taken together, these findings suggest, for the first time, that the PRC2-dependent epigenetic remodeling may contribute to the bacteria-induced mucosal imprint to sensitize recurrent diseases.

Heightened inflammatory response to bladder reinfections has been previously observed and largely attributed to the adaptive immunity(49–51). It is therefore a surprise to find that the urothelial cells also contribute to the heighten inflammatory response. Intriguingly, the PRC2-dependent inflammatory response in the urothelial progenitor cells appears to be at the expense of terminal differentiation of superficial cells. Loss of the superficial cell layer grants the invading pathogens access to deeper bladder tissue and predisposes the host to chronic cystitis(29). On the one hand, the UTI-induced immunological memory in urothelial cells accelerates bacterial clearance by a heightened inflammatory response; on the other hand, it increases the risk of chronic infection due to the impaired urothelial homeostasis. UPEC have evolved multiple strategies to subvert the host’s defense(64), including intracellular localization(26, 27, 29), filamentation(65), attenuation of cytokine production(66–68), killing of natural killer cells(69), limiting antigen presentation(50), and promoting cell survival(63). Our findings here provide the initial evidence suggesting that UPEC have also taken advantage of epigenetic remodeling of the bladder urothelial cells to regulate host innate immunity to benefit the pathogens.

In the repeated infection model, *Eed^cKO^* mice appear to have partial resistance to reinfections while maintaining urothelial tissue homeostasis (Figure 4). While the underlying mechanism remains to be defined, we have noted that the aryl hydrocarbon receptor (AhR) signaling pathway is significantly upregulated in *Eed* mutant urothelium (Figure 1). AhR is a ligand-dependent transcription factor that is able to detect bacterial pigments and regulate antibacterial defense(70). AhR regulates expression of several LPS-responsive genes(71), and mitigates inflammatory response to primary LPS challenge(72). On LPS rechallenge, however, the AhR-dependent mechanism promotes an establishment of a disease tolerance state that resists bacterial infection and preserves tissue homeostasis(72). These observations suggest an alternative mechanism through which targeting PRC2 may mitigate symptoms of UTIs by improving bladder urothelial homeostasis.

Injuries awaken the quiescent bladder urothelium, causing urothelial progenitor cell proliferation and differentiation to repair the damage(8). UTI-induced *Ezh2* expression coincides with reactivation of the cell cycle of urothelial progenitor cells. The basal cell proliferation rate is significantly reduced in the urothelium-specific *Eed* mutants during acute, repeated and superinfections, consistent with the previous observation that *Ezh2* expression is closely associated with cell proliferation during bladder urothelial development(9). Pathway analysis of the differentially expressed genes in the bladder urothelium between *Eed^cKO^* mice and controls suggest that activity of the *p53* tumor suppression pathway is significantly elevated in *Eed^cKO^* mice. Consistent with these observations, conditional knockout of *Kdm6a* in the bladder urothelial cells, a histone demethylase that catalyzes an opposing biochemical reaction of PRC2 methyltransferase, results in a significant decrease in the *p53* tumor suppressor pathway activity(73). Moreover, loss-of-function of *Kdm6a* significantly increases bladder cancer risk(73). Thus, the *PRC2/Kdm6a-p53* axis regulates urothelial progenitor cell proliferation and tumorigenesis. Collectively, the PRC2-dependent epigenetic program may play a dual role in urothelial progenitor cells – it enhances inflammatory host response through the *NF-**κ**B* signaling pathway and, at the same time, controls urothelial regeneration through the *p53* tumor suppressor pathway by promoting proliferation of basal progenitor cells and inhibiting differentiation of superficial cells.

In summary, the findings here suggest that UTI induces the PRC2-dependent epigenetic program in the bladder urothelium, which exaggerates inflammation and proliferation at the expense of differentiation and tissue homeostasis, and further suggest an non-antibiotic strategy to ameliorate UTIs by inhibiting the methyltransferase activity of PRC2. This study also raises several questions regarding mucosal imprint in the bladder. Future studies are needed to examine whether the immune and urothelial cells may share the similar or divergent roles to modulate the immunity; whether and how the UTI-induced epigenetic reprograming can be reversed to reduce the risk of bladder diseases; and whether targeting *Ezh2* enzymatic activity may reduce the risk and/or mitigate UTIs.

## Materials and Methods

Mouse lines including *Eed^F/F^, UpkII^Cre^* and *Shh*^Cre^ have been described previously in detail(9). CAG-TRAP mice were kindly provided by Dr. William T Pu, Boston Children’s Hospital and available at The Jackson Laboratory (Jackson Laboratory, 022367). C57BL/6J mice (000664) were obtained from The Jackson Laboratory; And C3H/HeNCrl mice (025) were obtained from Charles River. DZNep (1.5mg/kg, Selleckchem, S7120), EPZ005687 (2.5mg/kg, APExBIO, A4171) and cyclophosphamide (CPP, 300mg/kg, Sigma C7397) were administered to mice via intraperitoneal (i.p.) injection. Female mice between 6-8 weeks of age were used in the bladder infections as previously described(44). The uropathogenic *Escherichia coli* UTI89 strain was generously provided by Dr. Scott Hultgren (Washington University, St. Louis, USA). For the C^+^UTI superinfection model, mice received a single dose of cyclophosphamide (CPP, 300mg/kg, Sigma C7397, i.p.) 1 day prior to UTI89 inoculation. All animal studies were performed according to protocols reviewed and approved by the Institutional Animal Care and Use Committee at Boston Children’s Hospital.

## Author contributions

XL, ZB and CG designed the research studies. CG, MZ, XS, ZB and SZ conducted experiments and acquired data. XL, CG and MZ analyzed data. XL wrote the manuscript with the help from CG.

## Acknowledgments

We thank all members of the X.L. lab, particularly Drs. Satoshi Kaneko, Chao Yang and Christa Lam for technical supports and helpful discussions. We appreciate Drs. Rosalyn Adam and Eric Greer for critical and insightful comments on the manuscripts. This work was supported by NIH/NIDDK (1R01DK091645-01A1, X.L.), NIH/NCI (R21CA249701, X.L) and NHLBI (1R01HL136921, X.L.).

## Supplemental Materials and Methods

### Bladder infections

Female mice between 6-8 weeks of age were used in the bladder infections as previously described(Hung et al., 2009). The uropathogenic *Escherichia coli* UTI89 strain was generously provided by Dr. Scott Hultgren (Washington University, St. Louis, USA). Under isoflurane, 10^8^ colony forming units (CFU, in 50μl PBS) of UTI89 stationary culture were transurethrally inoculated into the bladder. Urine bacterial burdens were determined by serially diluting urine samples and counting CFU. For the repeated infections, infected mice were re-infected (up to 7 times) after 7 days of convalescence. For the C^+^UTI superinfection model, mice received a single dose of cyclophosphamide (CPP, 300mg/kg, Sigma C7397, i.p.) 1 day prior to UTI89 inoculation.

### Histology, inflammation score and immunofluorescent staining

Histology and immunostaining were performed as previously described(Guo et al., 2017). To quantify infiltration of neutrophils, we used four bladders per experimental group. Three paraffin sections from each of the bladders were H&E stained for histological analysis. To count infiltrating neutrophils, 200X microscopic fields of each section were used to count neutrophils and total number of cells. Neutrophils were identified based on their characteristic multilobed nucleus morphology. Percent of neutrophils of total cells was calculated and analyzed using Student’s t-test and p<0.05 was considered significant. Inflammatory score were determined using the described criteria(Hopkins et al., 1998). Two investigators were Antibodies used for immunostaining included Ezh2 (1:100 Cell signaling, 5246), H3K27me3 (1:100 Millipore 07-449), Ki67 (1:100 Abcam 16667), pHH3 (1:100 Millipore, 06-570), Krt5 (1:500 Abcam, ab53121; 1:500, Covance, SIG-3475), Upk3a (1:200 provided by Dr. T.T. Sun), Krt20 (1:50, Dako, M7019), I**κ**Bα (1:100 Cell signaling, 4814) and p65 (1:100, Santa Cruz, sc8008). Images were captured using a ZEISS fluorescence microscope.

### Chromatin immunoprecipitation assay (ChIP)

Chromatin immunoprecipitation was performed as previously described(Guo et al., 2017). Briefly, micro-dissected urothelial cells from 20 bladders per group were crosslinked with 1% formaldehyde at room temperature for 15 minutes. Cells were then lysed and sonicated to an average size of 500-1000 base pairs. Immunoprecipitation of chromatin lysates was performed using 1μg per reaction of the following antibodies: anti-H3K27me3 (Millipore, 07-449), anti-H3K4me3 (Motif Active, 39159), or control IgG. Immunoprecipitates were collected with Protein G Dynabeads (Invitrogen) and protein/DNA crosslinks were reversed with 5M NaCl. DNA was purified and ChIP DNA was analyzed by quantitative PCR using the following *Iκßa* locus-specific primers, −10kb region: forward, 5’TAA AGC ACA TGT GTG GGG TCT C; reverse, 5’ATC GAA CAT AGG GCC ATG AGA G; −0.3kb region: forward, 5’CAT CGG AGA AAC TCC CTG CG; reverse, 5’CAA ACC AAA ATC GCC CGG TG. Each sample was normalized using input DNA.

### RNA isolation, TRAP, RNA-seq and RT-qPCR

RNeasy RNA isolation kit (Qiagen) was used to isolate total RNA from micro-dissected bladder urothelium. Briefly, mouse bladder was harvested and placed immediately in cold autoclaved phosphate buffered saline (PBS). Under stereo dissecting microscope, the bladder was bisected to expose the lumen. With one forceps holding and stabilizing the bladder at the muscular layer, the urothelium was gently peeled off using another forceps. This process usually took less than two minutes. Genomic DNA was removed using gDNA Eliminator spin columns (Qiagen). For Translating Ribosome Affinity Purification (TRAP) isolation, mutant (*Shh^Cre/+^; Eed^f/f^; TRAP^f/f^*) and control (*Shh^Cre/+^; Eef^f/+^; TRAP^f/f^*) mouse bladders at embryonic 18.5 (E18.5) were micro-dissected and stored in −80°C after ten minutes of incubation with 100μM cycloheximide buffer. Pooled bladder samples (100mg each) were used for TRAP RNA purification as described(Zhou et al., 2013). Illumina 2500 was used for RNA-seq (20M, PE150). Cutoff for differentially expressed genes (DEGs) was based on *padj* < 0.05. Pathway analysis was done using the Ingenuity Pathway Analysis (IPA) software package (Qiagen). For quantitative real-time PCR (qRT-PCR) experiments, mRNA was reverse-transcribed into cDNA using the SuperScript^®^ III Reverse Transcriptase Kit (Invitrogen). qRT-PCR analyses were performed using SYBR Green (Roche) on an ABI-7500 detector (Applied BioSystems). Relative gene expression levels were normalized to the internal control *Gapdh*. The following gene-specific primers were used:

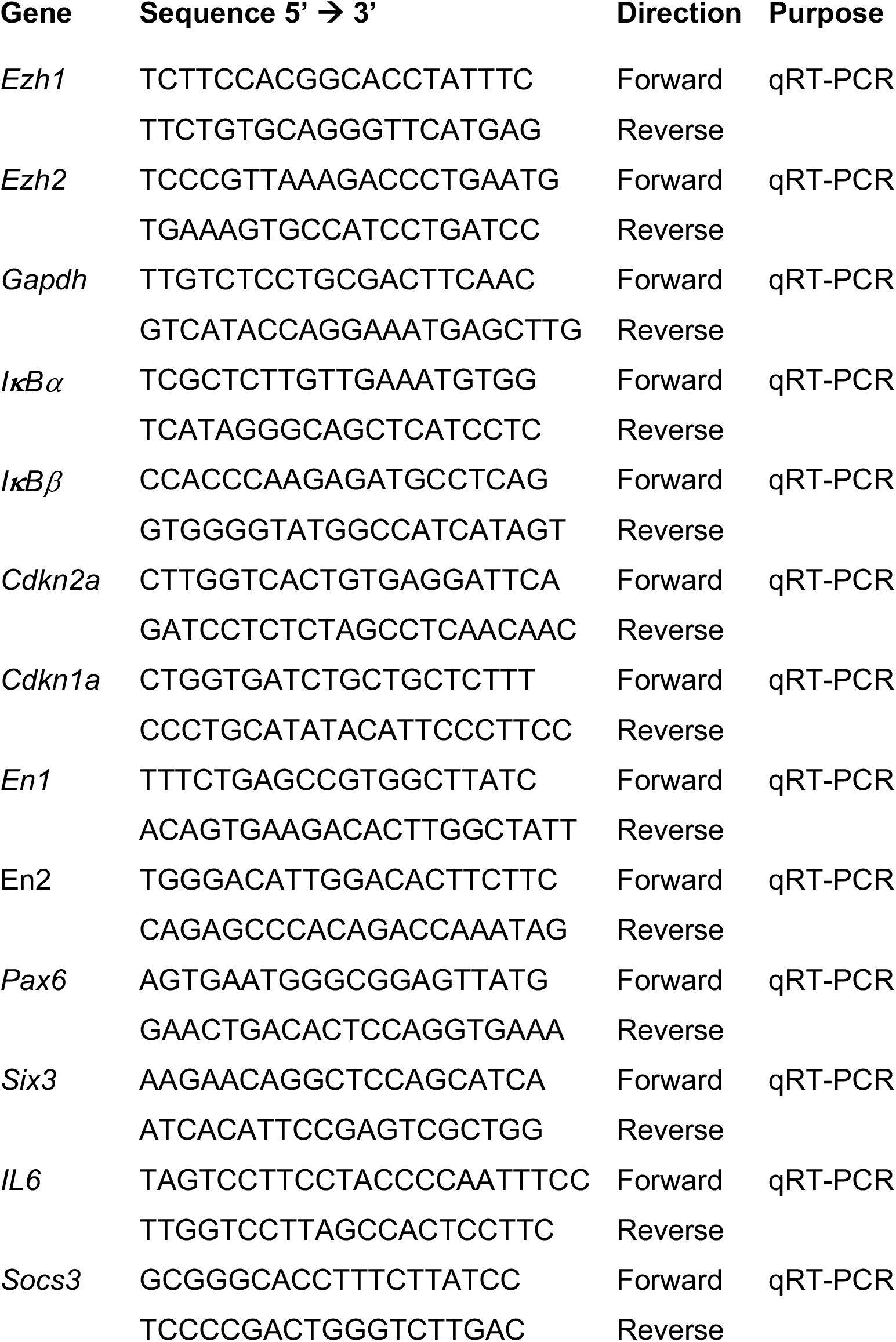
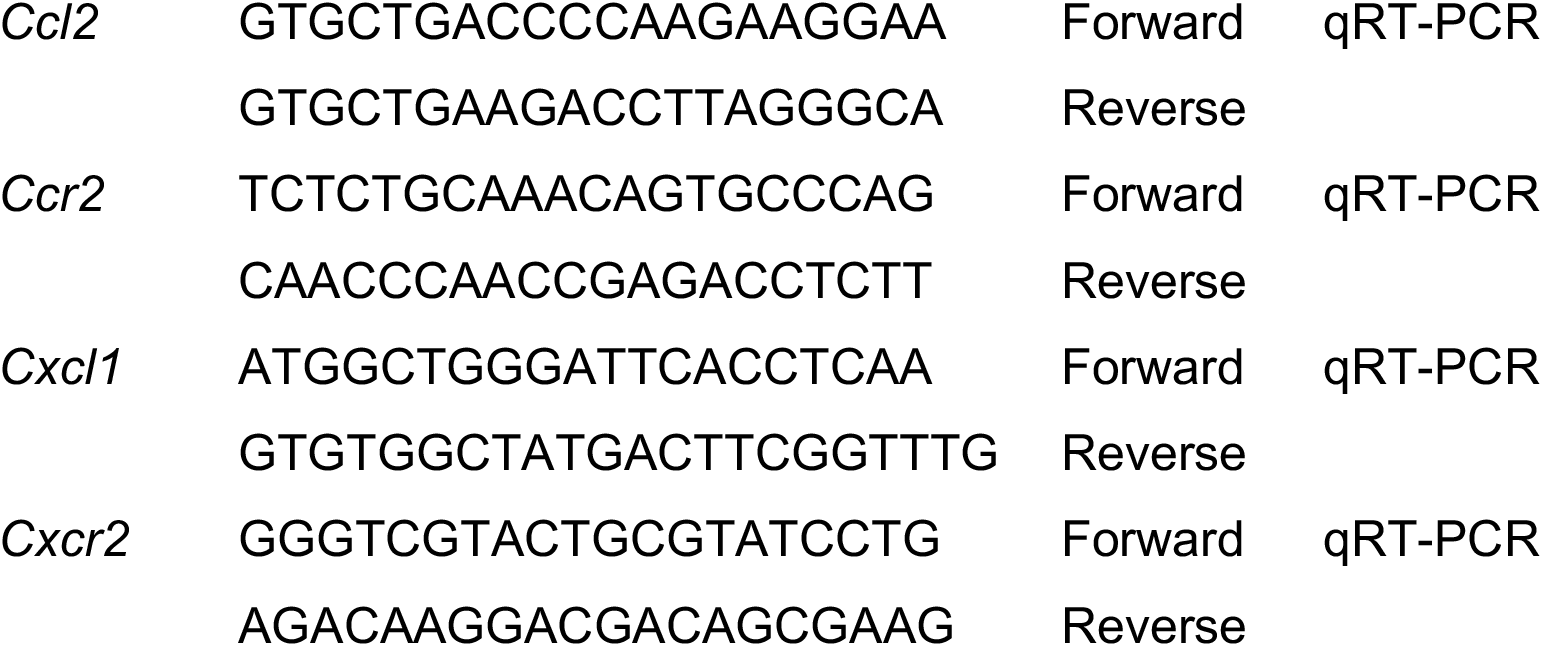

**Figure S1.**
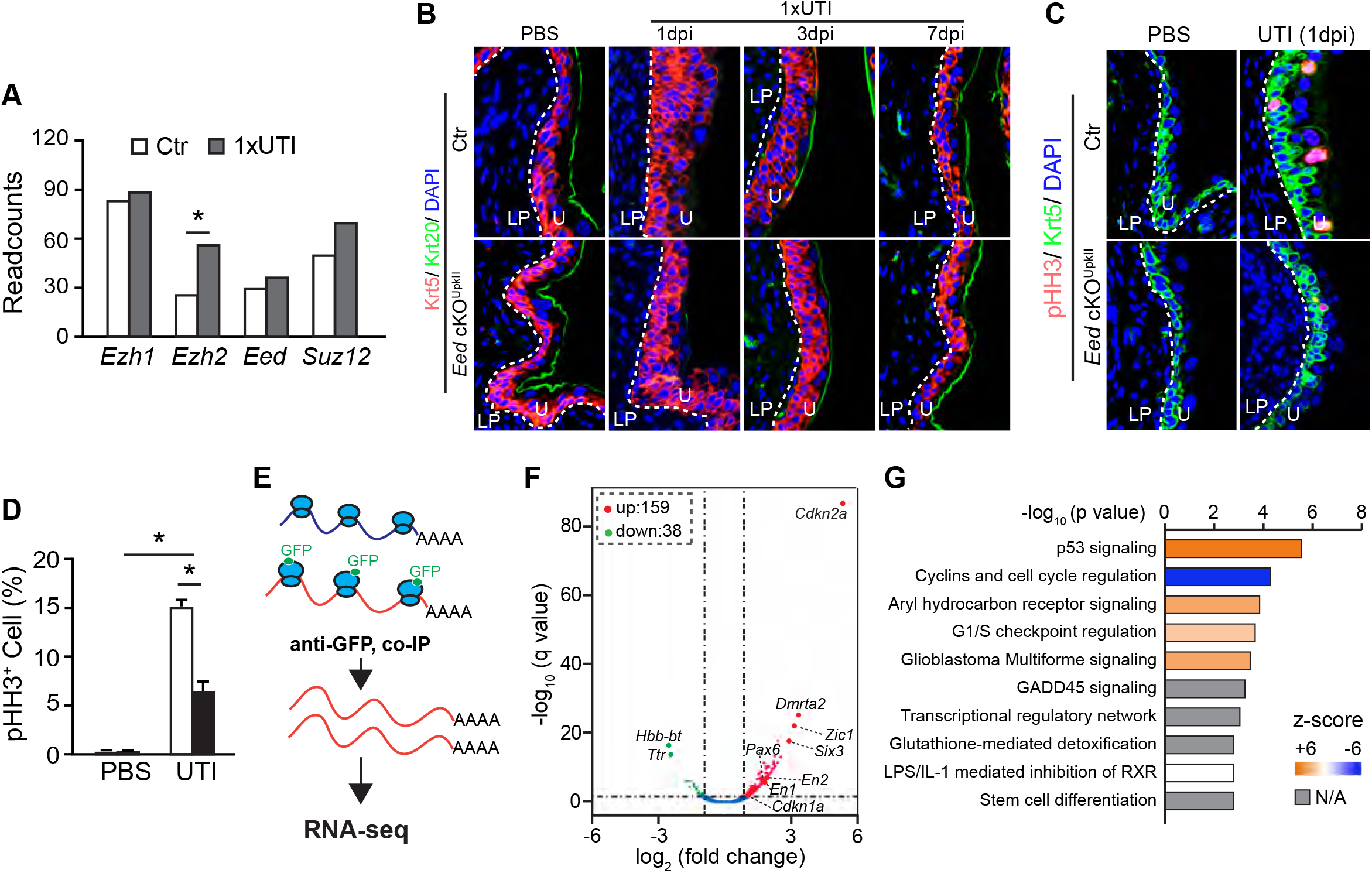
PRC2 controls urothelial progenitor cell proliferation during infection and development. (A) RNA-seq normalized read counts of *Ezh1, Ezh2, Eed* and *Suz12* mRNAs in adult urothelium at 1dpi. \**p*<0.05, Student’s *t*-test, n=2. (B) Krt5 (red) and Krt20 (green) immunostaining of the bladder sections of *Eed^cKO^* mice after an acute infection at 1 day-post-inoculation (dpi). Ctr, littermate heterozygous; PBS, vehicle treatment. Blue, DPAI counter staining; LP, lamina propria; U, urothelium. (C) Immunostaining of Krt5 (green) and pHH3 (red) of the bladder sections of *Eed^cKO^* mice after an acute infection at 1dpi. (D) Percentage of pHH3^+^ proliferating cells in the bladder urothelium relative to total cells. \**p*<0.05, Student’s *t*-test, n=3. (E) Schematic diagram of Translating Ribosome Affinity Purification (TRAP) strategy to isolate urothelial cell-specific TRAP RNAs for RNA-seq (Illumina 2500, 20M, pair end 150). mRNAs that were associated with polyribosomes were co-immunoprecipitated by GFP antibody, which recognizes the GFP-L10A fusion protein. The resulting TRAP RNAs were subjected to standard RNA-seq analysis. (F) Volcano plot of the differentially expressed genes (DEGs) in *Eed^cKO^* urothelium at embryonic day 18.5 (E18.5). n=2, *padj*<0.05. (G) The top 10 signaling pathways that are dysregulated in *Eed^cKO^* urothelium based on the Ingenuity Pathway Analysis (IPA) of DEGs shown in F. The *p53* molecular pathway is significantly activated.

**Figure S2.**
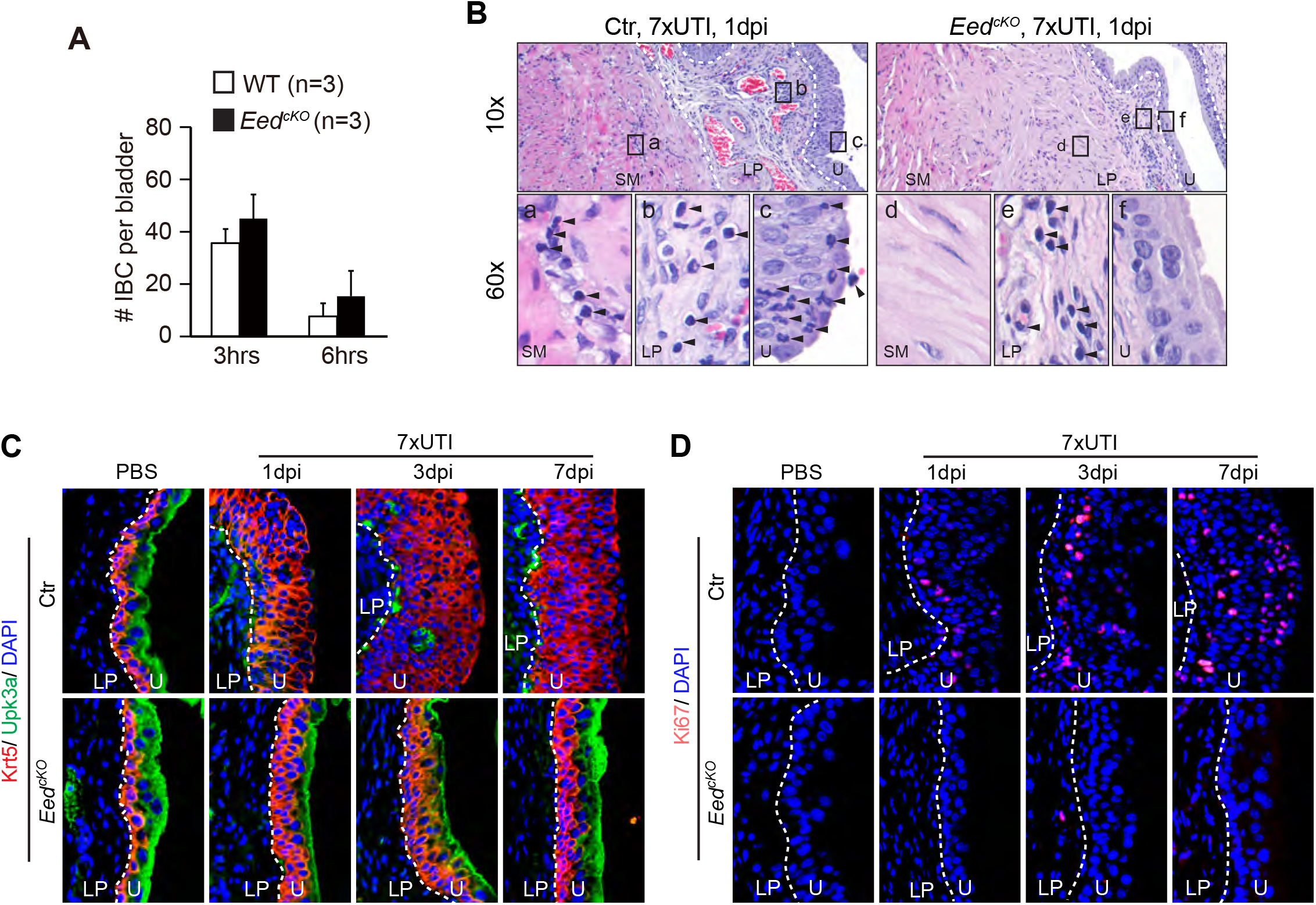
Genetic inactivation of PRC2 in the bladder urothelium improves outcomes of the repeated infections. (A) A number of intracellular bacterial communities (IBCs) observed in each bladder with 1xUTI at 3 or 6 hours post-inoculation. (B) Representative histological images of bladder sections from mice that were inoculated 7 times (7xUTI) and analyzed at 1 day-post-inoculation (dpi) of the last inoculation. LP, lamina propria; SM, smooth muscle; U, urothelium; dashed lines separate SM, LP and U; arrowheads, polymorphonuclear leukocytes. (C) Krt5 (red) and Upk3a (green) immunostaining of the bladder urothelium of mice with 7xUTI at 1, 3 and 7dpi. Blue, DAPI counter staining. (D) Ki67 immunostaining (red) of adjacent sections shown in C.

**Figure S3.**
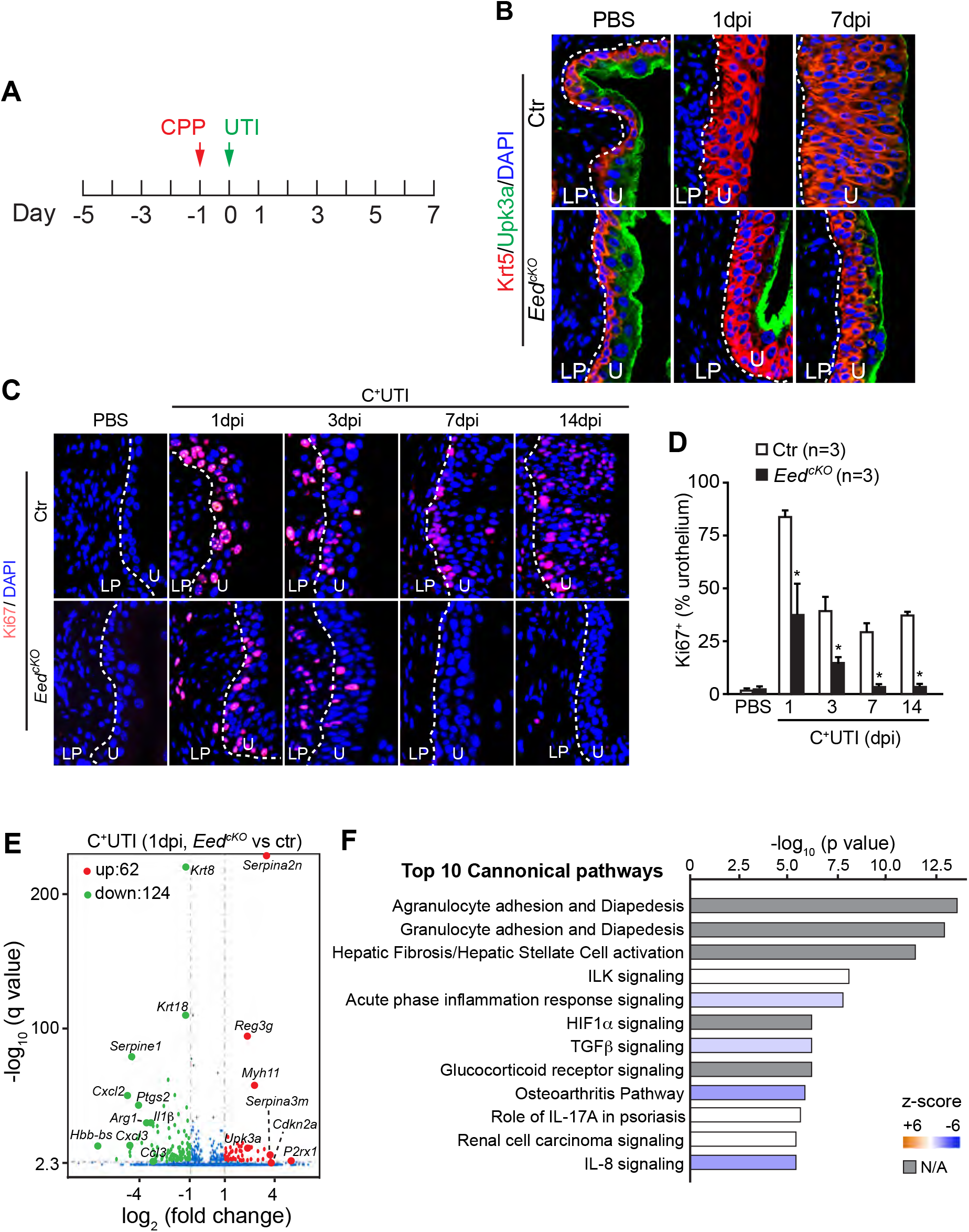
Genetic inactivation of PRC2 in the bladder urothelium improves outcomes of the C^+^UTI superinfection. (A) Schematic diagram of the C^+^UTI superinfection regimen. (B) Krt5 (red) and Upk3a (green) immunostaining of the bladder urothelium of mice that were C^+^UTI superinfected and analyzed at 1 and 7dpi. PBS, phosphate-buffer saline inoculated mice at 7dpi were used as controls. (C and D) Adjacent sections shown in A were stained with Ki67 antibody (red). Percentage of Ki67^+^ cells were quantitatively analyzed and shown in D. *p*<0.05, Student’s *t*-test, n≥3. (E) Volcano plot of the differentially expressed genes (DEGs) between *Eed^cKO^* and control urothelium at 1dpi of the C^+^UTI superinfection. n=2, *padj*<0.05. (F) Top ten canonical pathways identified by Ingenuity Pathway Analysis (IPA) of the DEGs that are affected in *Eed^cKO^* urothelium at 1dpi of the C^+^UTI superinfection.

**Figure S4.**
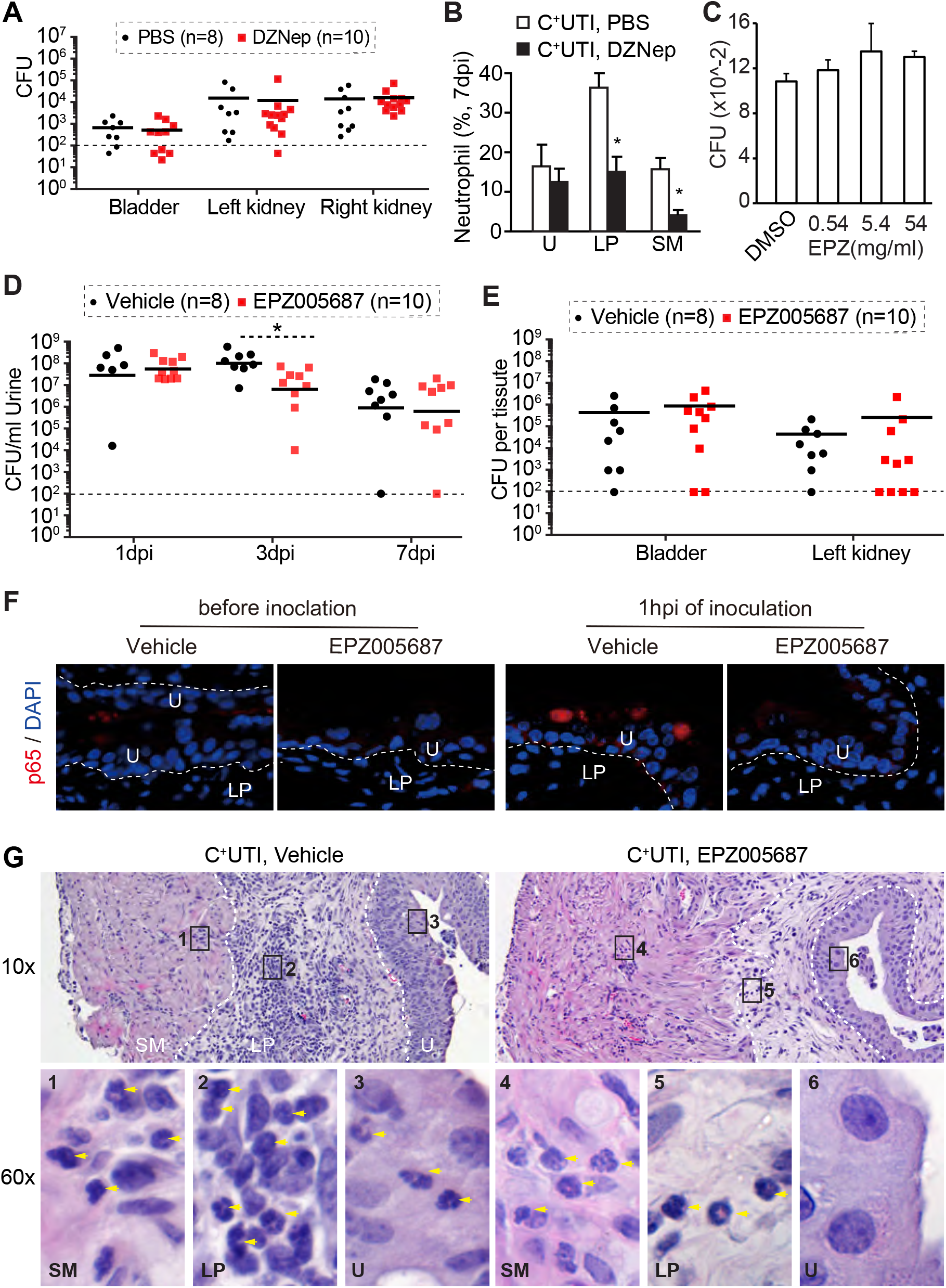
Ezh2 small molecule inhibitors improve outcomes of the C^+^UTI superinfection. (A) Bacterial counts in the bladder and kidneys of C57Bl/6 mice after the C^+^UTI superinfection with or without DZNep treatment. (B) Percentage of neutrophils found in the bladder of C57Bl/6 mice at 10dpi of the C^+^UTI superinfection. Treatments with DZNep and PBS vehicle control are indicated. *p*<0.05, Student *t*-test, n=3. (C) EPZ005678 has no detectable effect on UTI89 survival or proliferation *in vivo*. n=6. (D) Urine bacterial counts of C57Bl/6 mice after the C^+^UTI superinfection with or without EPZ005678 treatment. \**p*<0.05, Mann-Whitney test. (E) Bacterial counts in the bladder and kidneys of C57Bl/6 mice after the C^+^UTI superinfection with or without EPZ005678 treatment. (F) p65 immunostaining (red) the bladder sections that are treated with EPZ005678 or vehicle controls. Blue, DAPI counter staining. Dash lines separates the bladder urothelial and lamina propria layers. (G) Representative histological images of bladder sections of wild type C57Bl/6 mice at 7dpi of the C^+^UTI superinfections. Treatments with EPZ005687 and vehicle control are indicated. LP, lamina propria; U, urothelium; SM, smooth muscle; dashed lines separate SM, U and LP layers; arrowheads, polymorphonuclear leukocytes.

**Figure S5.**
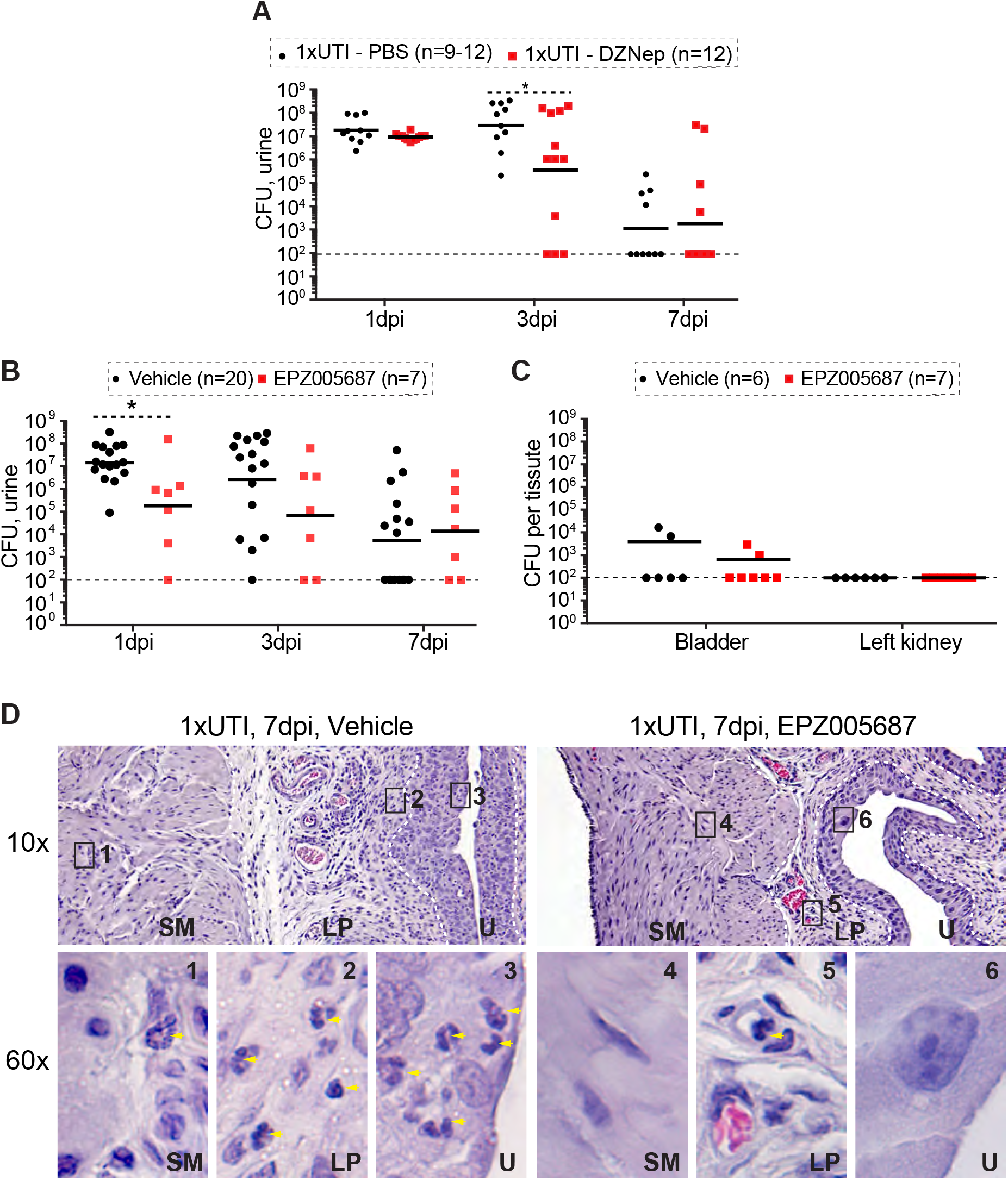
Ezh2 small molecule inhibitors improve outcomes of the chronic cystitis. (A and B) Urine bacterial counts of C3H/HeN mice after an acute infection after DZNep (A) or EPZ005687 (B) treatments. \**p*<0.05, Mann-Whitney test. (C) Bacterial counts in the bladder and kidney. (D) Representative histological images of bladder sections of wild type C3H/HeN mice at 7dpi. Treatments with EPZ005687 and vehicle control are indicated. LP, lamina propria; U, urothelium; SM, smooth muscle; dashed lines separate SM, U and LP layers; arrowheads, polymorphonuclear leukocytes.

